# Engineering Versatile Two-Dimensional Nanobody-Origami Architectures for Enhanced Antiviral Activity

**DOI:** 10.1101/2025.06.27.662057

**Authors:** Tingjie Song, Jazmin Galván Achi, Varada Anirudhan, Linh T.P. Le, Mehzabin Morshed, Xiaojing Wang, Lijun Rong, Xing Wang

**Affiliations:** Carl R. Woese Institute for Genomic Biology, University of Illinois at Urbana-Champaign, Urbana, IL 61801, USA; Holonyak Micro and Nanotechnology Lab, Grainger College of Engineering, University of Illinois at Urbana-Champaign, Urbana, IL 61801, USA; Department of Chemistry, University of Illinois at Urbana-Champaign, Urbana, IL 61801, USA; Department of Microbiology and Immunology, College of Medicine, University of Illinois at Chicago, Chicago, IL 60612, USA; Department of Bioengineering, Grainger College of Engineering, University of Illinois at Urbana-Champaign, Urbana, IL 61801, USA; VinUni-Illinois Smart Health Center, VinUniversity, Hanoi, Vietnam; Cancer Center at Illinois, University of Illinois at Urbana-Champaign, Urbana, IL 61801, USA

**Author notes:** Corresponding author: Xing Wang. T. S., J. G. A., and V. A. contribute equally to this work.

## Abstract

Considering the serious global health burden posed by pathogenic viruses, the development of effective antiviral molecules and therapeutic strategies is critical for improving human health. Designing and synthesizing a versatile and biocompatible molecular platform offering neutralization of both coronaviruses and retroviruses, two virus families that are greatly problematic to the human population, remains a significant challenge. Here, we report a programmable and versatile platform built on a two-dimensional (2D) DNA origami that enables nanoscale spatial control of multivalent nanobody (Nb) patterns for a broad-spectrum antiviral application. By site-selectively conjugating a Nb to a DNA oligonucleotide and placing multiple of them to predefined sites on 2D DNA origami, we synthesized hybrid nano-architectures with tunable Nb patterns designed to approximately match the geometric presentation of viral surface proteins. We demonstrate that such specific Nb spatial configurations significantly enhance both viral binding affinity and neutralization potency. For SARS-CoV-2, a coronavirus, a triangular Nb pattern with matched spacing with the spike proteins achieved an IC_50_ of 1.52 nM, representing a 171-fold improvement over monomeric Nbs. Extending this strategy to a retroviral virus, Human Immunodeficiency Virus (HIV) by utilizing a gp120 spike-binding Nb, we observed a 233-fold increase in neutralization efficiency using a patterned 2D Nb nano-architecture. These findings suggest a generalizable and versatile platform strategy for engineering potent antiviral agents through spatially optimized Nb presentation for a corresponding viral pathogen, offering a promising avenue for future antibody and Nb-based drug development.

---

The treatment and prevention of viral infections remain critical global health priorities due to their potential to cause serious illnesses and transmit rapidly. The global spread of the SARS-CoV-2 virus has led to over 7 million confirmed deaths,^1^ while approximately 39.9 million people were living with HIV as of 2023.^2^ These viral threats have placed immense strain on public health systems, driving increased investment and research. As a result, there is a need to develop effective antiviral therapeutics against highly impactful and pathogenic viruses like SARS-CoV-2 and HIV. Built on mechanistic insights into viral life cycles,^3, 4^ various strategies have been developed to disrupt viral replication, including entry inhibitors, reverse transcriptase inhibitors, integrase inhibitors, and protease inhibitors.^5-7^ Among these, entry inhibitors,^7^ which prevent viruses from entering host cells, offer a particularly direct approach to blocking infection at an early stage. However, designing a versatile platform strategy that can potently inhibit multiple important virus classes, including coronaviruses and retroviruses, remain a challenge.

The development of nanobody (Nb)-based therapies has emerged as a promising alternative among various strategies for combating viral infections,^8-10^ owing to Nb’s much smaller size, higher stability, and strong affinity for specific viral epitopes.^11^ For effective viral neutralization, a multivalent design with a precise spatial arrangement of multiple binders is critical.^12-18^ Such a multivalent design increases the number of potential binding sites across multiple epitopes, thereby enhancing the blockade of viral entry pathways and impairing the virus’ ability to infect host cells. Additionally, spatially organized Nb arrays amplify viral binding avidity,^19, 20^ where simultaneous interactions between multiple Nbs and their corresponding epitopes produce a much stronger and prolonged binding than monovalent interactions. Such multivalent engagement not only increases the durability of the antiviral effect but also improves resilience against viral mutations by enabling the targeting of conserved structural regions.^12, 21^ Thus, designing Nb therapies with well-defined spatial configurations holds great potential for developing robust, potent, and long-lasting antiviral treatments capable of addressing both existing and emerging viral threats.

In our previous study, we employed gold nanoparticles (AuNP) as a core carrier to present Nbs for blocking SARS-CoV-2 infections.^21^ However, the performance of the strategy can be further improved by overcoming the following key limitations brought by the AuNP-based platform. First, spherical AuNP exhibits minimal cytotoxicity in biological systems;^22^ Second, it is challenging to accurately control the spatial arrangement of surface-displayed Nbs surrounding the spherical AuNPs.^23, 24^ Additionally, we aim to develop a platform strategy with more versatility demonstrating efficacy against other pathogens, such as the retrovirus human immunodeficiency virus (HIV).

Herein, we present a versatile platform strategy for neutralizing different infectious viral pathogens with collectively higher Nb potency, offering both enhanced biocompatibility and precise Nb spatial controls. By leveraging the programmability and biocompatibility of DNA origami, we have arranged Nbs with high spatial precision to align with the spatial and geometric pattern of targeting viral surface proteins. To demonstrate the platform’s broad-spectrum potential, we validated its neutralization efficacy against both SARS-CoV-2 (a coronavirus) and HIV-1 (a retrovirus) as a showcase using respective virus-targeting Nbs (**Fig. 1**).

**Fig. 1.**
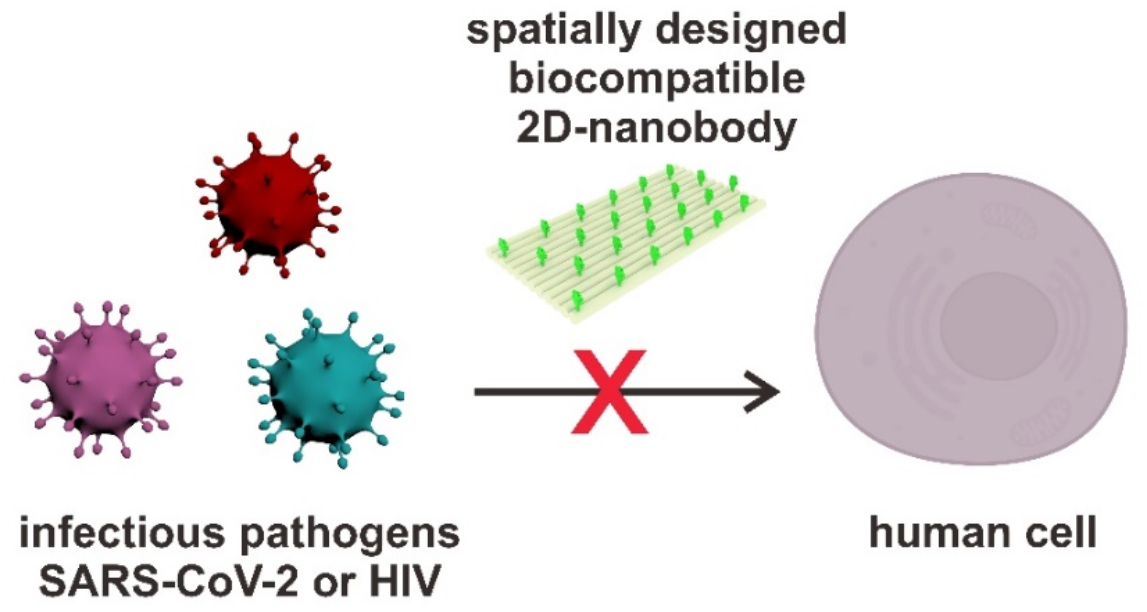
Schematic illustration of a versatile two-dimensional nano-architecture with spatially patterned Nbs for protecting human host cells from viral pathogen infections.

Shown in **Fig. 2a**, a two-dimensional (2D) nano-architecture designed and synthesized by using DNA origami technology^25-28^ (referred to as “2D origami” herein) was constructed by folding M13mp18 scaffold DNA with a set of staple strands with programmed sequences (**Table S1-3**) that include those with specific extensions for subsequent hybridization with DNA–nanobody conjugates. The assembled 2D origami nanostructures were purified using polyethylene glycol (PEG) precipitation.^29^ Atomic force microscopy (AFM) images reveal well-formed rectangular-shaped 2D origami with dimensions of approximately 90 nm in length and 70 nm in width. To achieve site-specific conjugation of native Nb to a DNA oligo, a two-step bioconjugation protocol was employed^21^ (**Fig. 2b**). First, an azide group was introduced at the N-terminus of the GlyHis_6_-tagged Nb via an acylation reaction.^30^ It was followed by a click chemistry reaction with a dibenzocyclooctyne (DBCO)-modified DNA oligo. Successful Nb– DNA conjugation was confirmed by native polyacrylamide gel electrophoresis (PAGE), which shows a clear shift in migration compared to the free DNA oligos, due to the increased molecular weight and reduced mobility imparted by the Nb-DNA conjugation (**Fig. 2c**).

**Fig. 2.**
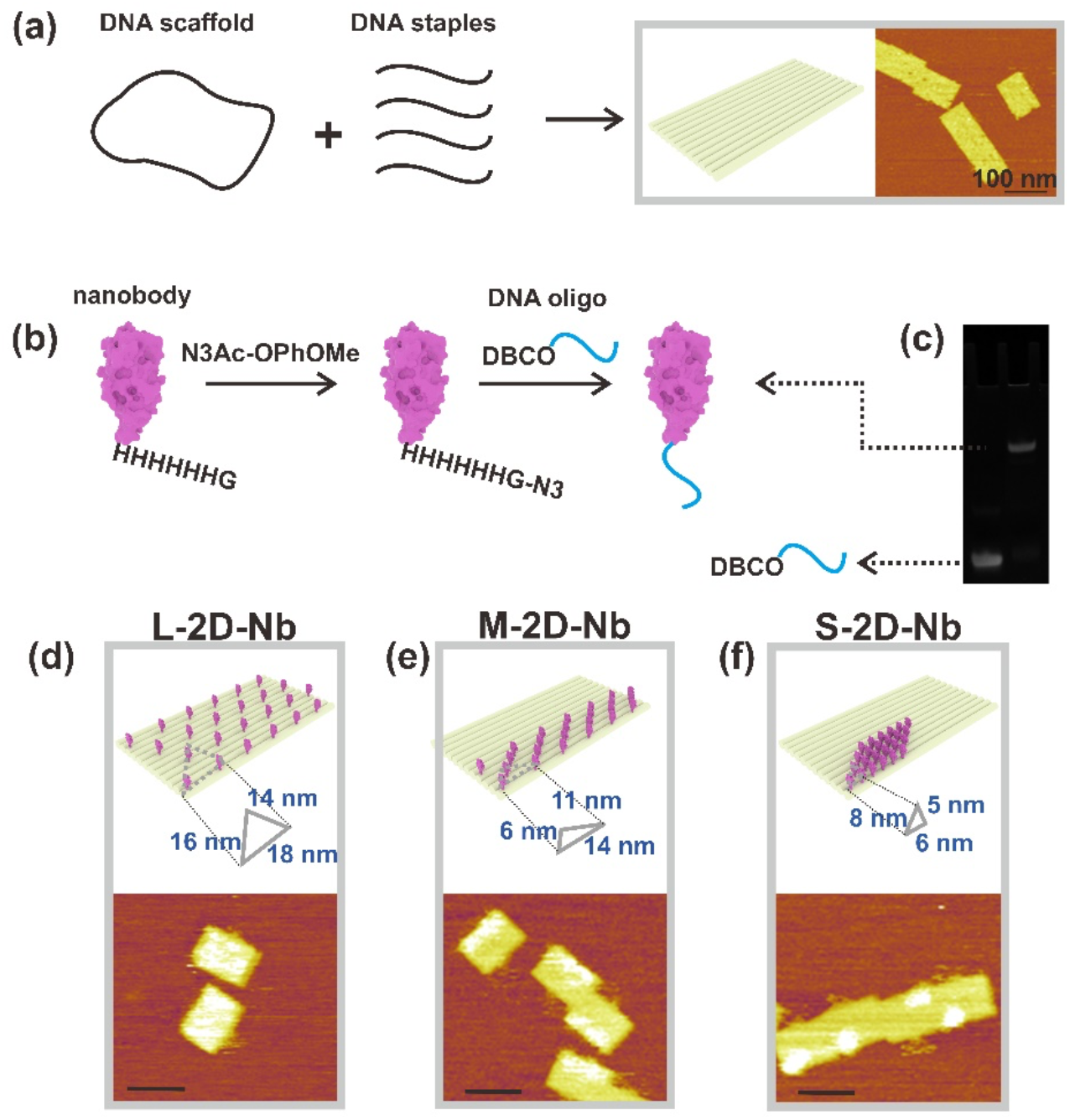
Design, synthesis, and characterization of multivalent Nb nano-architectures built on a 2D origami platform. (**a**) Assembly of a 2D DNA origami nanostructure. (**b**) Schematic of site-specific Nb-DNA conjugation. (**c**) PAGE gel analysis confirms successful Nb–DNA conjugation. Left lane: DBCO-DNA; right lane: Nb-DNA conjugate. (**d–f**) AFM images show different spatial arrangements of Nb on 2D DNA origami by design. The scale bars indicate 100 nm.

With both 2D origami and Nb–DNA conjugates available, we designed multivalent Nb patterns on a 2D origami platform with precise spatial arrangements. Given the trimeric nature of viral class-I fusion proteins displayed on both SARS-CoV-2 and HIV^31-34^, Nbs are arranged in triangular clusters to proximately match the geometry of viral spike proteins. Literature reports have estimated the center-to-center distance between monomers within a given class-I fusion trimeric protein structure to be approximately 7–8 nm^31-34^. To test the relationship between the accommodation of such intra-trimeric spacing and the outcome of Nb’s antiviral efficacy, we engineered various Nb patterns on 2D origami with programmable spatial arrangements using modified staple strands (**Table S1-3**). **Fig. 2d** shows three Nb-arrangements patterned on the 2D origami. In the large-spacing 2D Nb (L-2D-Nb) structure, 24 Nbs are placed across the entire 2D origami with the intra-tri-Nb spacing of 18, 16, and 14 nm, as indicated by white dotted regions under AFM imaging. In contrast, medium-spacing (M-2D-Nb) and small-spacing (S-2D-Nb) configurations concentrate the same number of Nbs as stripe-and cluster-like on the 2D origami, which include tighter spatial arrangements of 14– 11–6 nm and 8–6–5 nm, respectively.

To validate the hypothesis that spatially organized Nbs with an approximate matching pattern to that of viral trimeric spikes, we employed surface plasmon resonance (SPR) in quantifying the binding avidities between different Nb-DNA origami constructs (2D-Nbs) and the SARS-CoV-2 spikes. 2D DNA origami only (without Nbs) exhibits negligible binding with the spikes, confirming that the DNA origami platform alone does not contribute to protein target engagement (**Fig. S1**). After functionalizing with Nbs, the binding affinity of a 2D-Nb was found to be dependent on the spatial arrangement of Nbs on the 2D DNA origami platform (Fig. S2). Specifically, the S-SARS-Nb achieved the highest binding affinity, with an equilibrium dissociation constant (*K*_*D*_) of 0.36 pM, representing a 21-fold or 10-fold improvement over the

L-SARS-Nb or M-SARS-Nb. These results suggest that both Nb spacing and spatial distribution geometry influence multivalent binding strength. The spatial arrangement of S-SARS-Nb that better matches the pattern of trimeric spikes appears to promote cooperative Nbs-spikes binding for enhanced avidity, whereas suboptimal spacing offered by L-SARS-Nb or M-SARS-Nb reduces the binding strength.

We then evaluated the antiviral efficacy of the 2D multivalent patterned-Nb structures using a SARS-CoV-2 pseudovirus (PV) model that displays wild-type (WT) trimeric spikes (**Fig. 3**). In the absence of Nb treatment, the SARS-CoV-2 spikes bind to the ACE2 receptor on host cells (BHK-21-ACE2 cells used herein) for viral entry. When treated with 2D-Nbs, the SARS-CoV-2 surface-displayed spikes interact with multiple Nbs patterned on the 2D origami, forming tight virus–2D-Nb complexes that block virus-cell interactions. Such neutralization efficacy has been quantified by assessing the number of virions that invade host cells (**Fig. 3a**), and the half-maximal inhibitory concentration (IC_50_) has been determined for each of the three 2D-Nb constructs (L-SARS-Nb, M-SARS-Nb, S-SARS-Nb). As shown in **Fig. 3b-3d**, increasing concentrations of each 2D-Nb construct have led to dose-dependent inhibition of viral infection. The IC_50_ values were 5.20 nM for L-RBD-Nb, 2.89 nM for M-RBD-Nb, and 1.52 nM for S-RBD-Nb. Notably, the S-2D-Nb design exhibits a 171-fold enhancement in neutralization efficiency compared to the monomeric Nb with an IC_50_ of 260 nM (**Fig. S3**), highlighting the significant advantage of multivalent presentation in inhibiting virus infections. Additionally, cell viability assays confirmed the excellent biocompatibility of the 2D constructs (**Fig. S4**). Thus, the 2D-Nb constructs, integrating Nb specificity with the structural programmability of DNA origami, offer a versatile and biocompatible platform strategy for the development of advanced antiviral therapeutics.

**Fig. 3.**
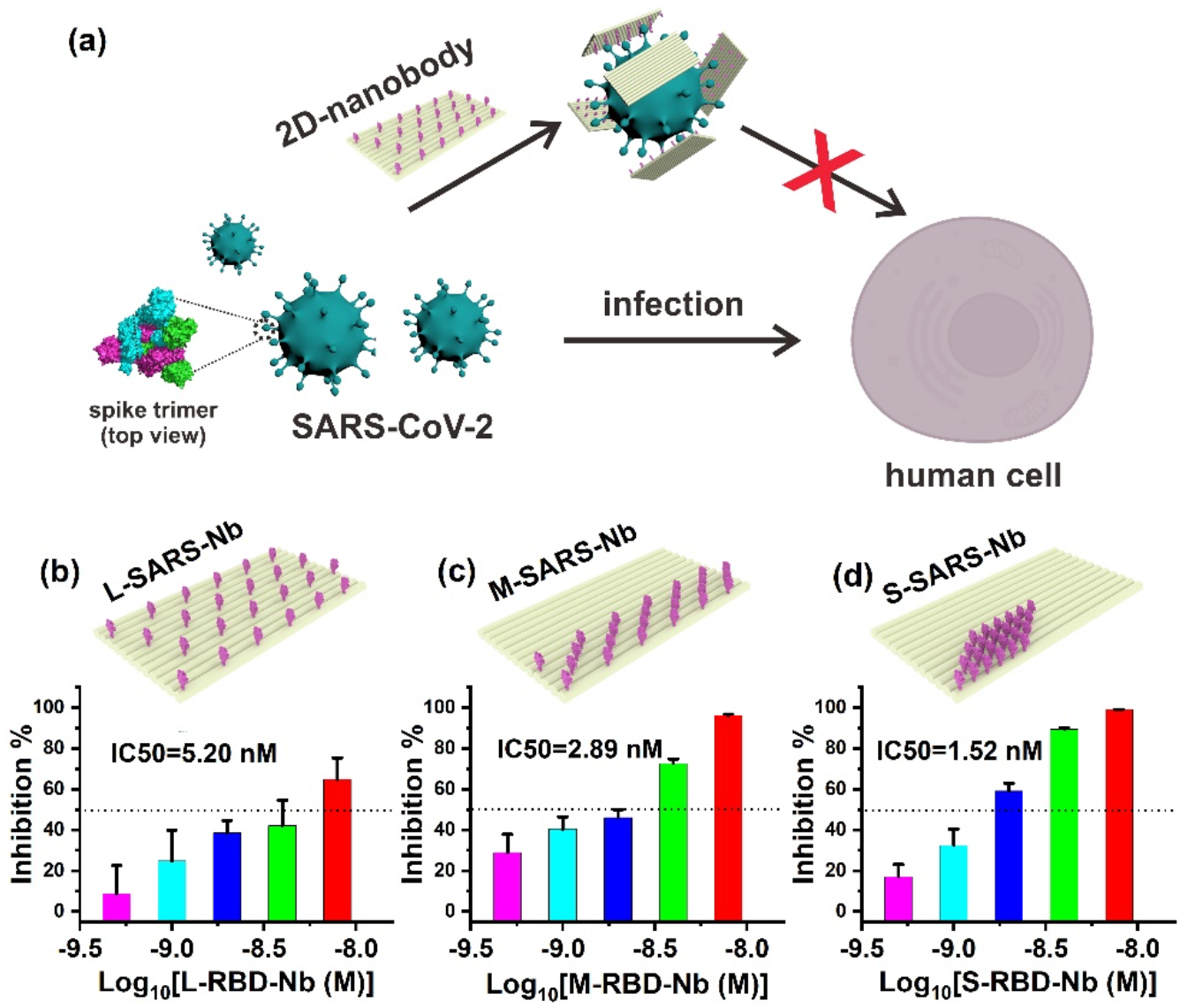
Neutralization of SARS-CoV-2 virus using 2D-Nb constructs. (**a**) Schematic illustration of the viral inhibition mechanism, where Nbs are organized on 2D DNA origami bind to the SARS-CoV-2 spikes with approximately matched spatial pattern and thus, preventing spike-ACE2 interactions. (**b–d**) Virus neutralization efficacy of respective 2D-Nb constructs with different spatial arrangements: (**b**) L-SARS-Nb, (**c**) M-SARS-Nb, and (**d**) S-SARS-Nb. Antiviral efficacy was quantified by IC_50_ values in SARS-CoV-2 virus infection assays.

Next, we test the possibility of utilizing our 2D-Nb platform strategy for the neutralization of HIV-1 infections, which remain a global health threat. The HIV-1 surface trimeric spike glycoprotein, GP120, is essential for viral entry by binding to CD4 receptors on host T lymphocytes. Structural studies^33, 34^ have revealed that GP120 forms a trimeric complex with an inter-subunit spacing of ∼ 8 nm (**Fig. 4a**), approximately matching the nanobody spacing patterned on one of our 2D-Nb constructs (8 nm–6 nm–5 nm). Based on the GP120 structural information, we functionalized the 2D origami with GP120-targeting Nbs to generate a spatially patterned Nb construct (termed 2D-HIV-Nb). AFM imaging confirms the successful formation of 2D-HIV-Nb constructs, with white dots indicating Nbs localize near one corner of the 2D origami (**Fig. 4b and Fig. S5**). When the HIV PVs incubated with 2D-HIV-Nb (**Fig. 4c**) at concentrations of 0.5 nM or above, viral infectivity was nearly abolished (**Fig. 4e**). The IC_50_ of 2D-HIV-Nb was determined to be 0.051 nM, representing a 233-fold improvement in inhibition efficiency compared to the monomeric Nb with an IC_50_ of 11.89 nM (**Fig. 4d**). As a negative control, unmodified 2D origami itself showed negligible viral neutralization, confirming the critical role of spatially arranged nanobodies in antiviral activity (**Fig. 4f**).

**Fig. 4.**
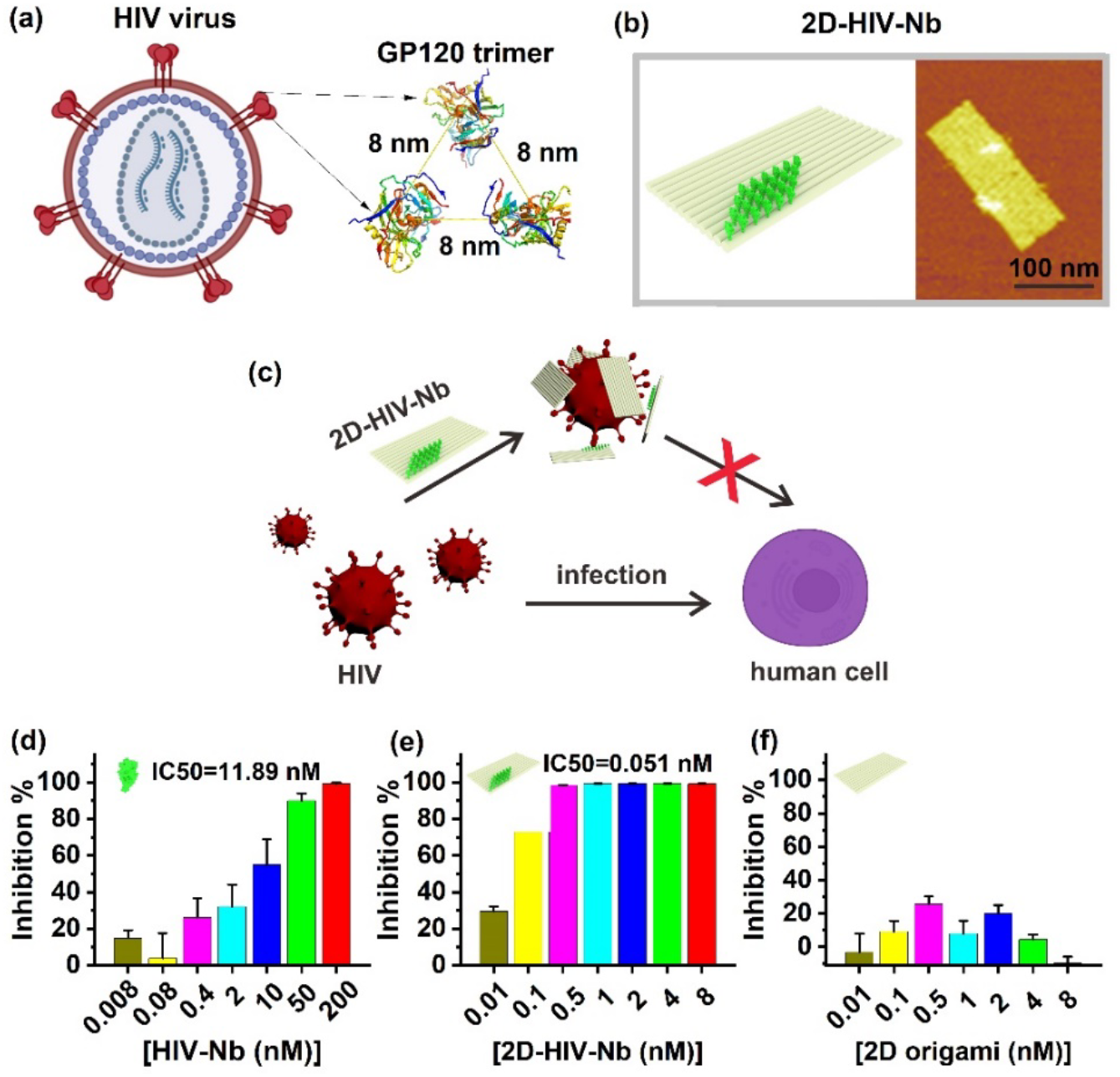
HIV-1 pseudovirus neutralization using 2D-Nb constructs. (**a**) Schematic of HIV-1 and the trimeric structure of its surface spike glycoprotein GP120. (**b**) AFM image confirms the spatially patterned Nbs on DNA origami (2D-HIV-Nb). (**c**) Schematic of the HIV pseudovirus neutralization assay, where 2D-HIV-Nb blocks the interaction between viral surface GP120 spikes and CD4 receptors on host cells. Comparison of neutralization efficiencies of (**d**) monomeric Nb, (**e**) 2D-HIV-Nb, and (**f**) 2D origami only control.

In summary, we have designed and synthesized three 2D Nb-DNA origami hybrid constructs for testing the relationship between Nb spatial patterns with their collective antiviral efficacies as showcased by using SARS-CoV-2 and HIV-1 models. 2D-Nb constructs were synthesized with controlled Nb spacing, achieved by hybridizing Nb–DNA conjugates onto extended staple strands of the 2D origami structure. The spatial arrangements of Nbs were confirmed using atomic force microscopy (AFM) imaging. Notably, different Nb arrangements yield different binding affinities and antiviral efficacies. For instance, a triangular configuration with 8 nm–6 nm–5 nm spacing (S-SARS-Nb) achieves the highest binding affinity to SARS-CoV-2 spike protein (*K*_*D*_= 0.36 pM) and corresponds to the most effective viral neutralization outcome (IC_50_ = 1.52 nM). Extending this strategy to HIV, we demonstrated that a 2D-HIV-Nb nanostructure with an approximate spatial matching with HIV-1 trimeric spike GP120 structure has improved neutralization potency by 233-fold compared to monomeric Nbs. These results highlight the importance of pattern recognition in enhancing multivalent Nbs-virus interaction in designing Nb-based antiviral strategies. The 2D-Nb platform provides a powerful and modular approach for advanced antibody and Nb-based drug design, with broad potential for combating different viruses and preparing for the next pandemic.

## Acknowledgements

The study was supported by grants from NIH R21AI166898 and R01AI159454.

